# Novel automaticity index reveals a cognitive ability-related decline in gait automaticity during dual-task walking

**DOI:** 10.1101/2023.07.31.551290

**Authors:** Shuqi Liu, Andrea L. Rosso, Emma M. Baillargeon, Andrea M. Weinstein, Caterina Rosano, Gelsy Torres-Oviedo

**Author notes:** **Correspondence:** Gelsy Torres-Oviedo.

## Abstract

Gait automaticity refers to the ability to walk with minimal recruitment of attentional networks typically mediated through the prefrontal cortex (PFC). Reduced gait automaticity is common with aging, contributing to an increased risk of falls and reduced quality of life. A common assessment of gait automaticity involves examining PFC activation using near-infrared spectroscopy (fNIRS) during dual-task (DT) paradigms, such as walking while performing a cognitive task. However, neither PFC activity nor task performance in isolation measures automaticity accurately. For example, greater PFC activation could be interpreted as worse gait automaticity when accompanied by poorer DT performance, but when accompanied by better DT performance, it could be seen as successful compensation. Thus, there is a need to incorporate behavioral performance and PFC measurements for a more comprehensive evaluation of gait automaticity. To address this need, we propose a novel automaticity index as an analytical approach that combines changes in PFC activity with changes in DT performance to quantify gait automaticity. We validated the index in 173 participants (≥65 y/o) who completed DTs with two levels of difficulty while PFC activation was recorded with fNIRS. The two DTs consisted of reciting every other letter of the alphabet while walking over either an even or uneven surface. We found that as DT difficulty increases, more participants showed the anticipated decrease in automaticity as measured by the novel index compared to PFC activation. Furthermore, when comparing across individuals, lower cognitive function related to worse automaticity index, but not PFC activation or DT performance. In sum, the proposed index better quantified the differences in automaticity between tasks and individuals by providing a unified measure of gait automaticity that includes both brain activation and performance. This new approach opens exciting possibilities to assess participant-specific deficits and compare rehabilitation outcomes from gait automaticity interventions.

## 1 Introduction

The ability to move around in the community is essential for independent living (Patla and Shumway-Cook, 1999; Rosso et al., 2013; Sheppard et al., 2013; Rantakokko et al., 2016). Successful community mobility requires gait automaticity (Clark et al., 2014; Van Swearingen and Studenski, 2014; Brustio et al., 2018), which refers to the automatic control of walking with minimal recruitment of attentional networks primarily residing in the prefrontal cortex (PFC) (Van Swearingen and Studenski, 2014; Clark, 2015). Gait automaticity declines with age (Gschwind et al., 2010; Montero-Odasso et al., 2011; Van Swearingen and Studenski, 2014; Clark, 2015) and may contribute to an increased risk of falls (Gschwind et al., 2010; Montero-Odasso et al., 2011). Therefore, it’s important to have a standardized way to quantify gait automaticity to help evaluate fall risks and promote mobility in older adults.

Gait automaticity can be evaluated by behavioral or neurophysiological assessments. The behavioral assessment uses dual-task walking (DT, i.e., walking and performing a cognitive task simultaneously) and compares the performance in DT relative to single-task (ST, i.e., either walking or performing a cognitive task) (Paul et al., 2005). High automaticity is reflected by similar walking and cognitive performance in DT and ST (Paul et al., 2005; Brauner et al., 2021; Longhurst et al., 2022). When walking is automatic, it is assumed to require minimal attentional resources resulting in no impact on the participant’s performance on the cognitive task (Clark, 2015). The neurophysiological approach more directly measures the attentional resources that are required to complete the task. The attentional resources are usually quantified by the cortical activation of the PFC (Holtzer et al., 2011; Clark, 2015; Herold et al., 2017; Menant et al., 2020) using noninvasive brain imaging technologies such as functional near-infrared spectroscopy (fNIRS).

DT paradigms integrated with fNIRS-based PFC measurements provide a useful avenue to assess gait automaticity. However, it is challenging to interpret PFC measurements and task performance independently during automaticity assessments. Specifically, a small change in PFC activation from ST to DT has opposing interpretations depending on task performance. Namely, a small increase in PFC activation alongside good task performance may indicate high automaticity (Clark, 2015). In contrast, when a small increase in PFC activation is coupled with poor task performance, it could indicate inefficiency in recruiting neural resources or the task is beyond capacity, as suggested by the CRUNCH model (Reuter-Lorenz & Cappell, 2008). Similarly, a large increase in PFC activation from ST to DT yields different interpretations contingent on task performance. A larger increase in PFC activation paired with maintained task performance could imply successful compensation from PFC for other brain regions whose structures and integrities have declined with aging, as suggested by several theories such as the HAROLD and STAC models (Cabeza, 2002; Reuter-Lorenz and Cappell, 2008; Park and Reuter-Lorenz, 2009; Maillet and Rajah, 2013; Fettrow et al., 2021). Conversely, when a large increase in PFC activation is paired with poor task performance, it could represent unsuccessful compensation or dedifferentiation, i.e., non-task-specific overactivation (Cabeza, 2002; Cabeza and Dennis, 2012; Festini et al., 2018; Fettrow et al., 2021). Therefore, there is a need to consider PFC activation and task performance simultaneously to characterize gait automaticity more accurately.

Given the interrelation of PFC activation and behavioral performance, analyzing them together may provide a more comprehensive measure of an individual’s gait automaticity than examining either alone. To address this need, we propose a novel automaticity index that combines PFC activity and DT performance to quantify gait automaticity. The objective of the study was to test the ability of the index to differentiate the change in automaticity 1) between tasks within the same participants, and 2) between participants. For objective 1, to compare between tasks, we computed the automaticity index in older participants (≥ 65 years of age, n=173) who completed two difficulty levels of DT. We expected worse automaticity index at greater levels of task difficulty. For objective 2, to compare between participants, we tested if individual’s cognitive abilities, measured by Mini-Mental State Exam scores, were related to automaticity. We anticipated that better cognitive ability would be correlated with better automaticity and that associations would be stronger for the automaticity index compared to either behavioral or neurophysiological measures alone.

## 2 Materials and methods

### 2.1 Participants

The data included participants from three previously published studies: Program to Improve Mobility in Aging, n = 42 (Brach et al., 2020); Neural Mechanisms of Community Mobility, n = 29 (Aizenstein et al., 2008); and Move Monongahela-Youghiogheny Healthy Aging Team, n =102 (Ganguli et al., 2010). The 3 datasets included similar experimental protocols but were collected for different purposes across two different lab spaces with two different fNIRS systems (same model) and with different experimenters.

All study participants were at least 65 years old, able to walk unassisted, and had no major neurological conditions. More details of the medical conditions and inclusion criteria were reported in the published studies. The Institutional Review Board at the University of Pittsburgh approved the studies, and all participants gave written informed consent.

#### 2.1.1 Participant Characteristics

In all studies, age, sex, race, and highest level of education were self-reported. General cognitive ability was assessed using the Mini-Mental State Exam (MMSE), a commonly used screening instrument for cognitive impairments (Folstein et al., 1975). For participants from the Program to Improve Mobility in Aging study, the Modified Mini-Mental State (3MS) Test was collected and the MMSE scores were derived from the 3MS scores by extracting only items equivalent to the MMSE and recalculating a score.

### 2.2 Experimental Paradigm

Participants performed single or dual tasks on an oval track with a 15-meter straight walkway on each side. One side had a standard surface (even) and the other side had wood prisms underneath carpets (uneven; Hoppes et al., 2020; Thies et al., 2005). All participants performed four experimental trials and each trial included a pseudo-randomized sequence of four conditions outlined here: two single tasks (ST) and two dual tasks (DT) (Fig. 1). The order was pseudo-randomized such that walking continued along the oval track. Single tasks included a motor single-task, walking on the even surface (walk), and a cognitive single-task, standing while reciting every other letter of the alphabet starting from B (standABC; Brandler et al., 2012; Holtzer et al., 2011; Verghese et al., 2012). The single tasks were used as a reference to compute the change in performance during DT to account for the different baseline abilities of each participant. The DT conditions required performing the cognitive task while walking on the even (evenABC) or uneven surfaces (unevenABC). The unevenABC condition was considered a harder DT condition than the evenABC because of the increased challenge of balancing and walking on the uneven floor surface.

**Figure 1.**
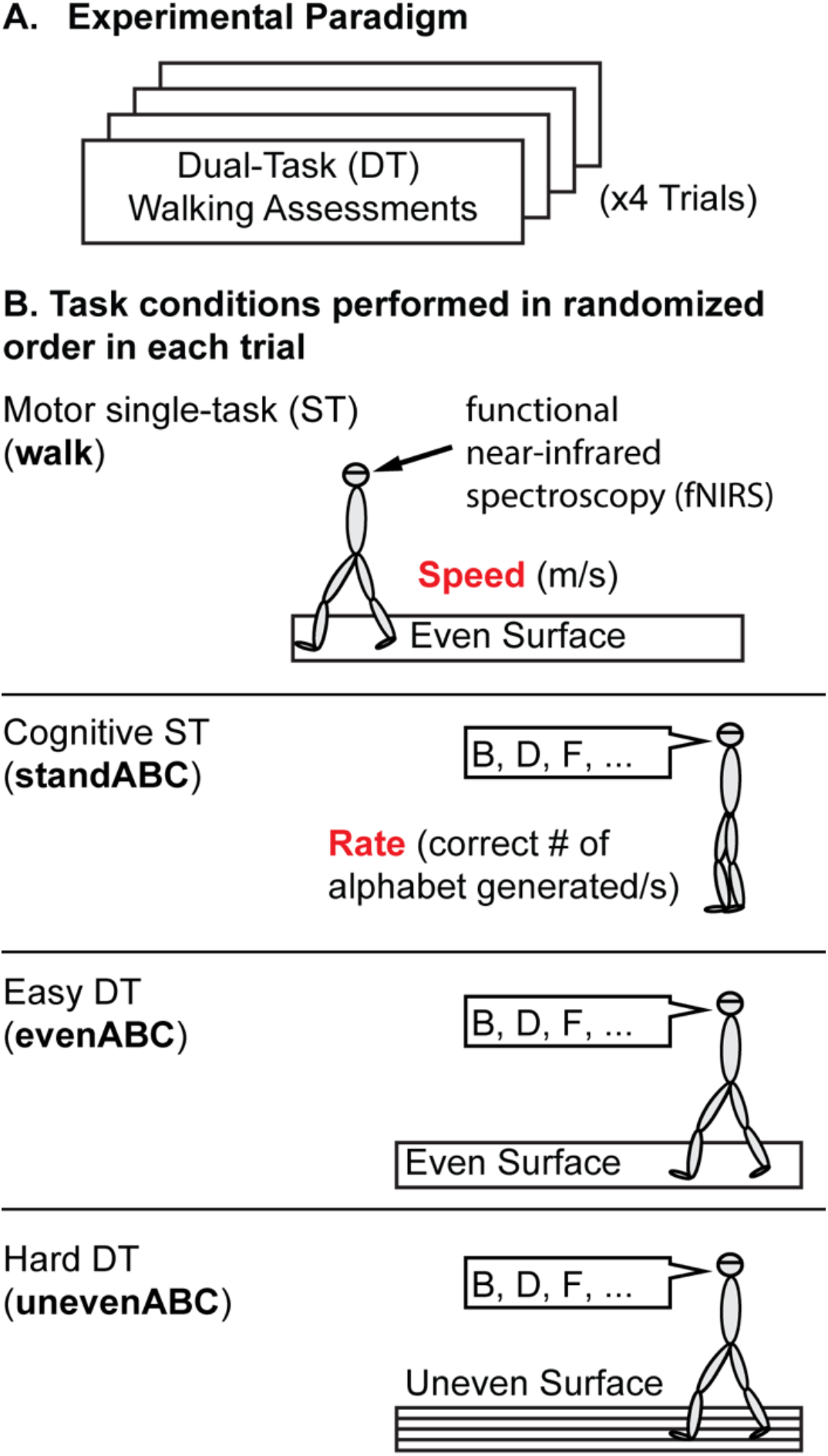
Experiment Protocol. **(A)** Overall protocol. Participants performed four trials of dual-task (DT) assessments. **(B)** Each trial includes four task conditions presented in pseudo-random order. The four tasks are: a motor single-task (ST): walking on an even surface, a cognitive ST: standing and reciting every other letter of the alphabet (standABC), and two DTs: reciting every other letter of the alphabet while walking on an even (evenABC) or uneven (unevenABC) surface. Motor performance is measured by gait speed (m/s) and cognitive performance is measured by the rate of correct letters generated. Prefrontal cortex activation was measured by functional near-infrared spectroscopy throughout the trial.

The task duration was 20 seconds for standABC. The duration for the motor ST (walk) and both DTs (evenABC and unevenABC) varied depending on the time the participant took to walk over the 15m straightway. Every condition was preceded by a quiet standing for 20 seconds, which was used as the baseline for fNIRS recordings. The quiet standing allowed the hemodynamic response to return to rest level such that relative changes in oxygenation of the blood could be computed for each task condition compared to quiet standing.

### 2.3 Data Collection

Motor performance was quantified by gait speed (m/s). Gait speed is computed as the distance (15m) divided by the time it took to walk the 15-meter walkway where time was measured by a stopwatch. Cognitive performance was quantified by the rate of correct letters of the alphabet generated per trial duration (correct letters/s). Average motor and cognitive performance for each condition across the four trials is reported.

PFC activation was measured using fNIRS which measures changes in blood oxygenation based on the distinct light absorption properties of oxygenated (Hbo) and deoxygenated (Hbr) hemoglobin (Miyai et al., 2001; Perrey, 2014; Hoppes et al., 2020). Participants wore an eight-channel continuous wave fNIRS headband (Octamon, Artinis Medical Systems, Netherlands) over their forehand during the entire experiment. The headband contained two detectors and eight sources covering both the left and right PFC regions. Near-infrared light transmitted at 840 nm and 760 nm was used to detect changes in Hbo and Hbr. Data was sampled at 10Hz and collected with the OxySoft software (Artinis Medical Systems, Netherlands).

### 2.4 fNIRS Data Analysis

fNIRS data was processed using the NIRS Brain AnalyzIR toolbox (Santosa et al., 2018) in MATLAB (Mathworks, Natick, Massachusetts). Light intensity was first converted to Hbo and Hbr measurements using the modified Beer-Lambert law with a partial path length factor of 0.1. A canonical model with the condition timing and duration was applied. The model was solved with an iteratively autoregressive pre-whitening least square approach to minimize motion artifacts. A student’s t-test was then performed on the regression coefficients and the t-score represents the changes in Hbo or Hbr in each task compared to baseline quiet standing. The results across four trials for the eight channels covering the whole PFC were combined using the toolbox to generate one ΔHbo and one ΔHbr for each participant for each task relative to quiet standing. Typically, an increase in PFC activity will be represented by an increase in oxygenated hemoglobin (i.e., a positive ΔHbo value) and a decrease in deoxygenated hemoglobin (i.e., a negative ΔHbr value).

ΔHbo often has a stronger signal-to-noise ratio (Menant et al., 2020) and therefore was used as the main measure of PFC activation for the remainder of the article. However, the same approach applies to both ΔHbo and ΔHbr measurements.

### 2.5 Automaticity index

The goal of the automaticity index is to create a single monotonic axis that combines performance and PFC activation to estimate automaticity where a larger index will always represent worse automaticity. Worse automaticity is reflected by more interference between dual tasks where they compete for the same attentional resources (Paul et al., 2005), leading to decreased task performance and increased recruitment of attentional resources (i.e., higher PFC activation).

The automaticity index combining cortical activation and performance is defined as the following:

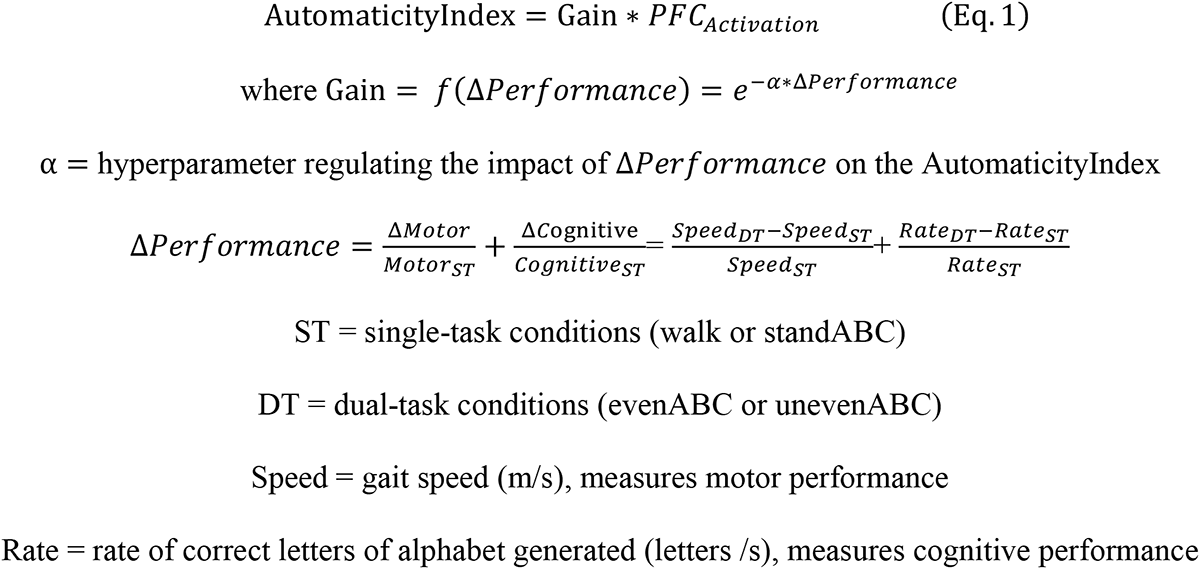

The automaticity index is generated by scaling PFC_Activation_ with a gain that is a function of the hyperparameter, α, and performance change, Δ*Performance*. Figure 2 demonstrates the process to compute the automaticity index using one of the datasets with three example participants highlighted to visualize the transformation of the data.

**Figure 2.**
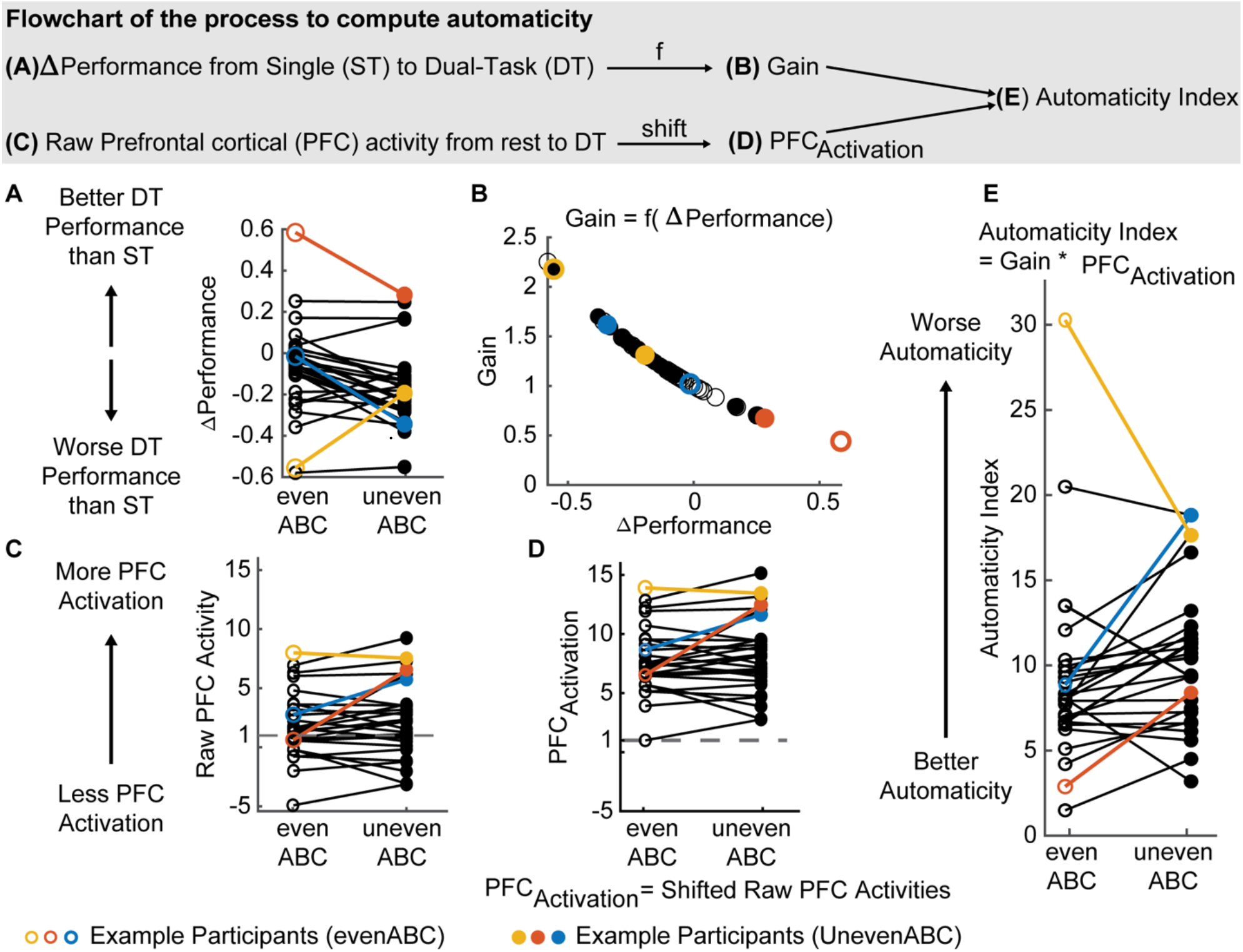
Schematics and flowchart illustrating the process to compute the automaticity index. The displayed example is one of the included datasets (n = 29) with 3 example participants highlighted (yellow, red, and blue). The evenABC (easier dual task, DT) task is represented by hollow circles and the unevenABC task is marked with filled circles. **(A)** Normalized performance change from single-task (ST) baseline to each DT. The performance combines both motor and cognitive performances. A positive value indicates performance is better in DT than ST. **(B)** Calculation of the gain from ΔPerformance. A large decrease in performance is mapped to a large gain (yellow empty circle). **(C)** The raw measure of PFC activity, which are t-scores representing changes in oxygenated hemoglobin (ΔHbo) concentration from rest to each DT. A positive value indicates increased Hbo concentration in DT compared to rest standing. A negative value indicates a decrease in Hbo concentrations in DT from rest. **(D)** The second term in the automaticity index equation, PFC_Activation_, simply shifts raw prefrontal cortex (PFC) activity values to be above or equal to 1. **(E)** The computed automaticity index, which is the multiplication of the gain **(B)** and the PFC Activation **(D)**.

The performance change, denoted as Δ*Performance* (Fig. 2A), is computed as the combined changes in cognitive and motor performances from ST to DT, normalized to performance in domain-specific ST (standABC and walk, respectively). The normalization accounts for individual differences in walking speed and alphabet performance at baseline. Cognitive and motor performances are weighted equally in Δ*Performance* to account for the different strategies employed by the participants during dual tasking since no specific prioritization instruction was given (Verghese et al., 2007; Yogev-Seligmann et al., 2010; Laguë-Beauvais et al., 2015). The definition of Δ*Performance* is similar to dual-task cost in existing literature (Brauner et al., 2021; Longhurst et al., 2022). A more negative Δ*Performance* value indicates performance was worse (i.e., participants walked slower and/or generated fewer correct alphabet letters) during DT compared to ST. Conversely, a positive Δ*Performance* represents better performance in DT compared to ST.

A negative sign was added before Δ*Performance* because we expected performance to decrease during DT (i.e., Δ*Performance* ≤ 0 ; Fig. 2A, majority of data points < 0). The negative Δ*Performance* reflects cognitive-motor interference during DT, which is expected in populations where gait automaticity is reduced, such as older adults.

The exponential weighting in the gain is chosen such that 1) performance decreases (Δ*Performance* < 0) scale the automaticity index up, with a large decrease weighted much more than a small decrease (Fig. 2B), 2) performance improvements (Δ*Performance* > 0, less common occurrences; Fig. 2B) scale the automaticity index down, but to a much smaller extent, and 3) unchanged performance will result in an index that is equal to the PFC activation. We weighted the gain in this way so that the index would more sensitively discriminate between participants with decreased rather than increased DT performance, as decreased performance with increasing task challenge is expected and is particularly relevant in the participant populations where gait automaticity is most often studied.

The hyperparameter *α* in the gain equation regulates the impact of Δ*Performance* on the automaticity index. *α* was optimized to minimize outliers while maximizing the sensitivity to reductions in gait automaticity as the task difficulty increased from the evenABC to unevenABC task. Specifically, the objective function is defined as:

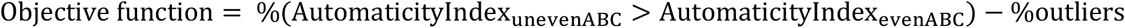

Outliers were defined as values more than three scaled median absolute deviations from the median, which is a robust measure of dispersion and outliers (Leys et al., 2013). The objective function value was evaluated for *α* ∈ [0, 5] with 0.1 increments. The optimal *α* was chosen as the smallest *α* that maximizes the objective function. At the start of the search range (*α* = 0), the gain from Δ*Performance* is equal to 1 and the automaticity index is equal to the PFC activation, which is the current standard in the field. The upper bound of the search range *α* = 5 was chosen heuristically for this dataset.

To maintain the monotonic property of the automaticity index and the relative differences across people and conditions, we turned all PFC_Activation_ values positive by shifting all PFC activity data, i.e., ΔHbo at DT compared to ST, by an offset (Fig. 2C, D), specifically:

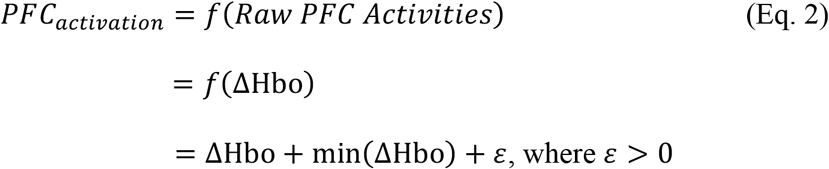

We shifted all PFC activation values by the minimum value across participants and tasks and then included an offset (*ε* = 1) such that the minimum PFC activation would be different than zero (Eq. 2). This was done to keep the automaticity index (the scaled PFC activation) monotonic while maintaining the relative difference in PFC activation across participants. Notice that Eq. 2 is a linear operation. Thus, the specific value of *ε* doesn’t impact the results. In this case, we chose *ε* = 1 for simplicity (Fig. 2D). Since the shift eliminates the sign of Hbo changes, our measure cannot determine how automaticity changes with respect to rest. However, it maintains the relative differences between tasks within a participant and across participants. In other words, the linear operation does not impact the goal of the automaticity index, which is to compare gait automaticity across tasks and individuals. A higher value before or after the shifting will always represent more PFC activation during the task.

If we were to use Hbr to represent the PFC activation, Hbr is first sign flipped to maintain the convention where higher *PFC*_*activation*_ always represents more PFC activity.

Figure 2 demonstrates the flowchart for computing the automaticity index using one of three included datasets. The blue participant is a typical example that behaves as we expected from the task design. As task difficulty increases, the blue participant decreased performance (Fig. 2A, negative slope) and increased PFC activation (Fig. 2C, D, positive slope), suggesting the harder task had more interference between walking and the alphabet tasks and required more attentional resources. In other words, the harder task was performed with less automaticity. As a result, the harder task had a larger automaticity index (Fig. 2E, positive slope). In contrast, the yellow participant increased performance (Fig. 2A) and decreased PFC activation (Fig. 2C, D) as the task became harder, suggesting the harder task was performed with improved automaticity (Fig. 2E, lower value in automaticity index for the harder task). This could result from the participant was not fully engaged in the easy task or the overall task design not being challenging enough.

The automaticity index is also effective in differentiating between participants. The red and blue participants had similar patterns of performance change (Fig. 2A, negative slopes for both) and PFC activation (Fig. 2C, D, positive slopes), but the red participant achieved better performance than blue (Δ*Performance* red > blue in both tasks). This observation suggests that with similar PFC resources, the red participant utilized the resources more effectively to achieve better performance. Consequently, the red participant should be considered to have better automaticity than the blue participant, i.e., lower automaticity index values (Fig. 2E, red much smaller than blue).

### 2.6 Statistical Analysis

For objective 1, to assess how each metric captured the anticipated task-difficulty changes in gait automaticity, we descriptively compared the automaticity index, PFC activation, and ΔPerformance by reporting the percentage of participants following the expected trend for each metric as task difficulty increases.

For objective 2, linear regressions were performed to test if the independent variable, MMSE scores, is related to the dependent variables, automaticity index, PFC activation, and performance respectively, while adjusting for demographics: age, sex, race, and highest level of education. Since the variables are on different scales, all variables were first standardized as Z-scores before the model fitting. Variance inflation factor (VIF) was computed to assess the multicollinearity among the independent variables in the adjusted models. Variables with VIF < 10 would be kept in the multiple regression models. Model significance was determined by F-test comparing the regression model with a constant model. To compare the impact of MMSE across the models, the standardized *β* is reported. Briefly, the standardized *β* represents in standard deviation unit how much a unit increase in MMSE will impact the dependent variable. We reported the ordinary R^2^ values, the p-values for the models, and standardized coefficient estimates with their respective standard deviations and p-values. One-sample Kolmogorov-Smirnov test was performed to check the normality of the standardized residuals.

Two participants did not follow the instruction to recite every other letter of the alphabet. A sensitivity analysis was included to test the impact of removing these two participants on the fitting of the models adjusted for demographics as well as on our conclusions. All analyses were performed in MATLAB and a statistical significance of *α* = 0.05 was used.

## 3 Results

Demographics, cognitive test results, cognitive and motor task performances, and fNIRS measurements of PFC activation in dual tasks are shown for the full sample in Table 1. Among the 22 non-white participants, one participant identified as Asian, one identified as American Indian or Alaskan Native, 19 identified as Black, and one identified as mixed or other race. Among the participants with education less than or equal to high school, only one participant’s highest level of education was less than high school.

**Table 1.** Demographic, clinical assessment results, task performances, and PFC activation (Hbo) of the study participants. All data are shown as mean±SD except for Sex, which is binary, Race, and Highest level of education, which are categorical.

| Variables | Mean (SD)<br>or n (%) |
| --- | --- |
| Sample size | 173 |
| Age (years) | 73.9 ± 5.6 |
| Female, n (%) | 108 (62.4%) |
| Race, n(%) |  |
| White | 151 (87.3%) |
| Non-white | 22 (12.7%) |
| Highest level of education, n (%) |  |
| Less than or equal to high school/equivalent | 43 (24.9%) |
| College | 87 (50.3%) |
| Postgraduate | 43 (24.9%) |
| Mini-Mental State Exam (MMSE) score (max 30) | 28.4 ± 1.8 |
| Gait speed at walk (motor single-task, m/s) | 0.97 ± 0.14 |
| Gait speed at evenABC (m/s) | 0.83 ± 0.17 |
| Gait speed at unevenABC (m/s) | 0.78 ± 0.16 |
| Rate of correct alphabet letters at standABC (cognitive single-task, letters/s) | 0.58 ± 0.14 |
| Rate of correct alphabet letters at evenABC (letters/s) | 0.62 ± 0.15 |
| Rate of correct alphabet letters at unevenABC (letters/s) | 0.59 ± 0.13 |
| Change in oxygenated hemoglobin (ΔHbo) from rest to evenABC (t-stats) | 1.88 ± 2.75 |
| ΔHbo from rest to unevenABC (t-stats) | 2.43 ± 3.22 |

### 3.1 Automaticity index increases as task difficulty increases

By experimental design, a harder task should require more attentional resources, i.e., increased PFC activation (Fig. 3A top left panel) and have poorer performance (Fig. 3B top left panel), resulting in worse automaticity, i.e., increased automaticity index (Fig. 3A and 3B bottom left panels).

**Figure 3.**
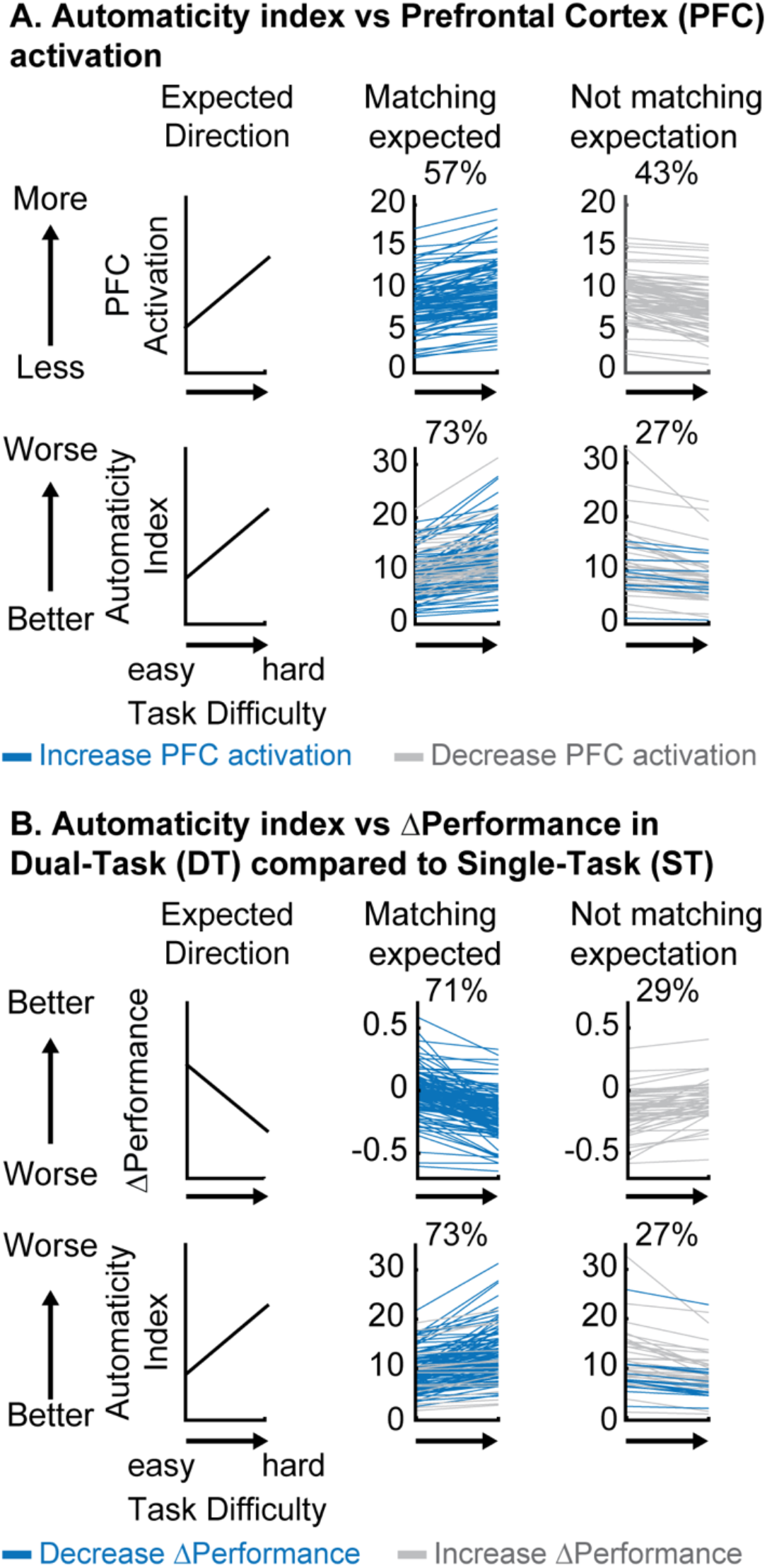
Descriptive comparison between automaticity index, PFC activation, and ΔPerformance as task difficulty increases. **(A)** Comparison between automaticity index and PFC activation at each dual-task (DT) relative to rest. Blue always represents participants who showed an expected increase in PFC activation (top middle panel) and grey always represents participants who decreased PFC activation (top right) as the task became more difficult. Colors in the bottom panel represent how participants from different groups at the top panel moved into different categories (increase or decrease) in automaticity index. Notice that some participants with an unexpected decrease in PFC activation will now have an expected increase in automaticity index after considering performance (grey in the top right moved to the bottom middle panel). **(B)** Comparison between automaticity index and ΔPerformance. Blue always represents participants who showed an expected decrease in Δ Performance as the task becomes harder (top middle panel) and grey always represents the ones who increased ΔPerformance (top right). Colors in the bottom panel represent how participants from different ΔPerformance at the top panel moved into different categories (increase or decrease) in automaticity index (bottom panel).

Only 57% of participants increased PFC activation as the task became harder (Fig. 3A top middle panel), showing no clear evidence of increased attentional control as task difficulty increased by this measure alone. However, 73% of participants increased automaticity index (i.e., had worse automaticity) as task difficulty increased (Fig. 3A bottom middle panel), matching our anticipated result that more challenging tasks would be performed with lower automaticity. Notice that the increase from 57 to 73% of participants following our expectation was primarily due to the grey participants whose PFC activation unexpectedly decreased with increased task difficulty (Fig. 3A top right). These grey participants recruited less PFC as the task became harder, but their performance declined, suggesting that PFC resources were not being used as effectively as needed by the task demand, which indicates reduced automaticity in the harder task.

In comparison, a larger percentage of participants (71%) decreased their performance (combined cognitive and motor performance) as the task became harder (Fig. 3B top middle panel), which was expected. A smaller percentage had better task performance during the more difficult task (29%, Fig. 3B top right panel). Some of these participants with improved task performance moved to the expected direction when using the automaticity index (grey in Fig. 3B bottom middle panel), perhaps identifying individuals who improved their performance at a cost of greater PFC recruitment. The increased PFC recruitment could represent a compensatory strategy to cope with the task demand, which correspond to reduced automaticity.

In summary, as task difficulty increases, more participants showed the expected decrease in automaticity as measured by the automaticity index and Δ Performance compared to PFC activations, suggesting that automaticity index and Δ Performance are more sensitive to the between task differences within the participants.

### 3.2 Greater automaticity index is related to worse cognitive function

All independent variables (MMSE score, age, sex, race, and highest level of education) had VIF less than 10 and therefore were all kept in the multiple regression models.

As expected, higher MMSE was associated with lower automaticity index at both tasks (Table2; evenABC: R^2^=0.06, p = 0.002; unevenABC: R^2^=0.05, p = 0.003) and this association at the easier task was robust to adjustment by age, sex, race, and highest level of education (Table 3; Figure 4; evenABC: R^2^=0.09, p = 0.02). The association between MMSE and automaticity index was slightly weaker in the adjusted model at the harder task(Table 3; Figure 4; unevenABC: R^2^=0.07, p = 0.06). Higher MMSE was also associated with lower PFC activation at both tasks (Table 2; evenABC: R^2^=0.03, p = 0.02, *β* = −0.18 ± 0.08; unevenABC: R^2^=0.02, p = 0.02, *β* = −0.13 ± 0.08), but these relations were either weaker (Table 3; evenABC: R^2^=0.07, p = 0.06, *β* = −0.13 ± 0.08) or not statistically significant (Table 3; unevenABC: R^2^=0.04, p = 0.40, *β* = −0.10 ± 0.08) after adjusting for covariates.

**Table 2.**
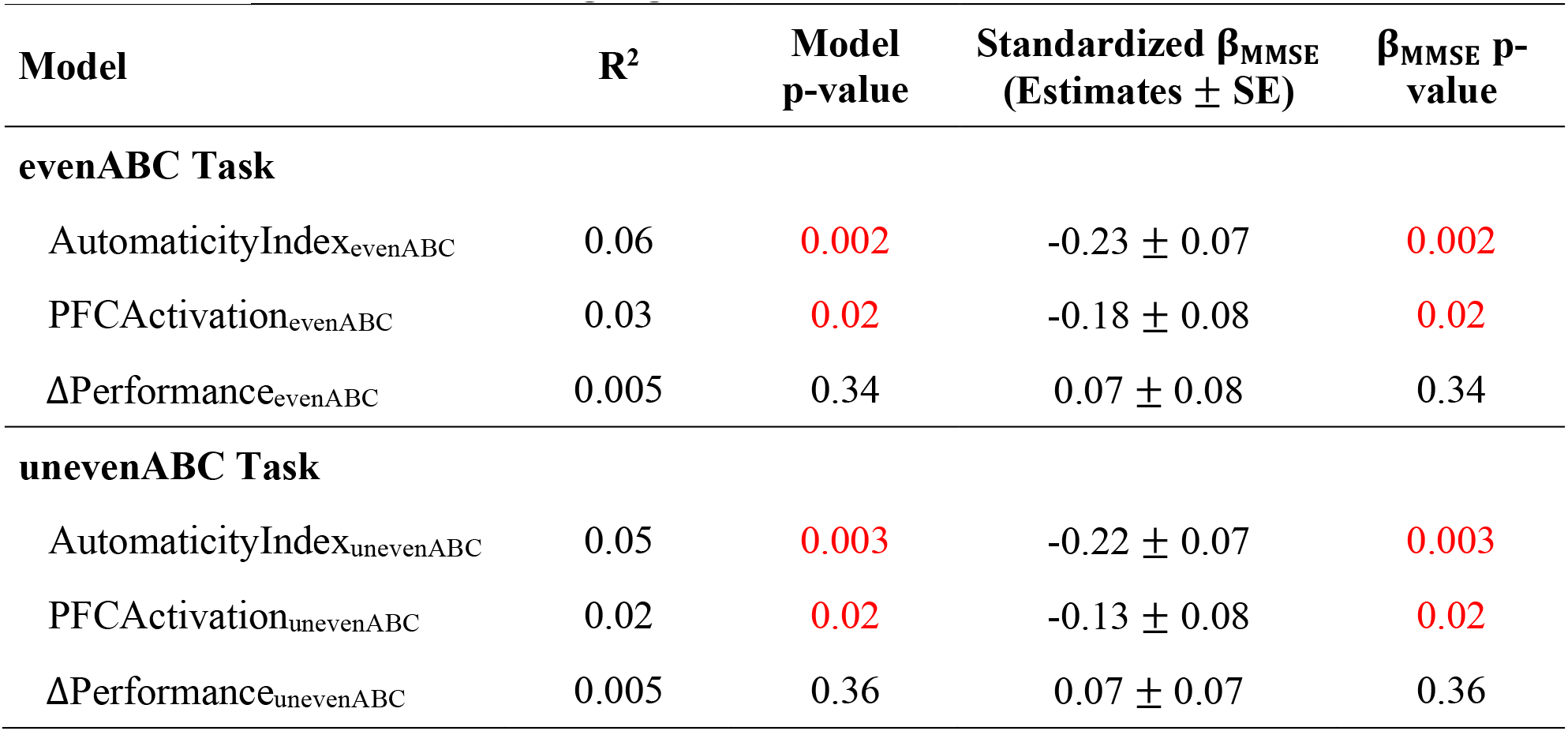
Unadjusted regression models of different gait automaticity measures with MMSE (n=173). Significant models are highlighted in red.

| Model | R<sup>2</sup> | Model p-value | Standardized $\beta_{\text{MMSE}}$<br>(Estimates ± SE) | $\beta_{\text{MMSE}}$ p-value |
| --- | --- | --- | --- | --- |
| **evenABC Task** |  |  |  |  |
| AutomaticityIndex<sub>evenABC</sub> | 0.06 | 0.002 | -0.23 ± 0.07 | 0.002 |
| PFCActivation<sub>evenABC</sub> | 0.03 | 0.02 | -0.18 ± 0.08 | 0.02 |
| ΔPerformance<sub>evenABC</sub> | 0.005 | 0.34 | 0.07 ± 0.08 | 0.34 |
| **unevenABC Task** |  |  |  |  |
| AutomaticityIndex<sub>unevenABC</sub> | 0.05 | 0.003 | -0.22 ± 0.07 | 0.003 |
| PFCActivation<sub>unevenABC</sub> | 0.02 | 0.02 | -0.13 ± 0.08 | 0.02 |
| ΔPerformance<sub>unevenABC</sub> | 0.005 | 0.36 | 0.07 ± 0.07 | 0.36 |

**Table 3.** Multivariable regression models of different metrics to quantify gait automaticity with MMSE (n=171) adjusted for age, sex, race, and highest level of education. Significant models are highlighted in red, and trending (p<0.1) models are highlighted in yellow.

| Model | R<sup>2</sup> | Model p-value | Standardized $\beta_{\text{MMSE}}$<br>(Estimates ± SE) | $\beta_{\text{MMSE}}$ p-value |
| --- | --- | --- | --- | --- |
| **evenABC Task** |  |  |  |  |
| AutomaticityIndex<sub>evenABC</sub> | 0.09 | 0.02 | -0.20 ± 0.08 | 0.01 |
| PFCActivation<sub>evenABC</sub> | 0.07 | 0.06 | -0.13 ± 0.08 | 0.10 |
| ΔPerformance<sub>evenABC</sub> | 0.02 | 0.77 | 0.08 ± 0.08 | 0.30 |
| **unevenABC Task** |  |  |  |  |
| AutomaticityIndex<sub>unevenABC</sub> | 0.07 | 0.06 | -0.19 ± 0.08 | 0.02 |
| PFCActivation<sub>unevenABC</sub> | 0.04 | 0.40 | -0.10 ± 0.08 | 0.21 |
| ΔPerformance<sub>unevenABC</sub> | 0.01 | 0.90 | 0.07 ± 0.08 | 0.41 |

**Figure 4.**
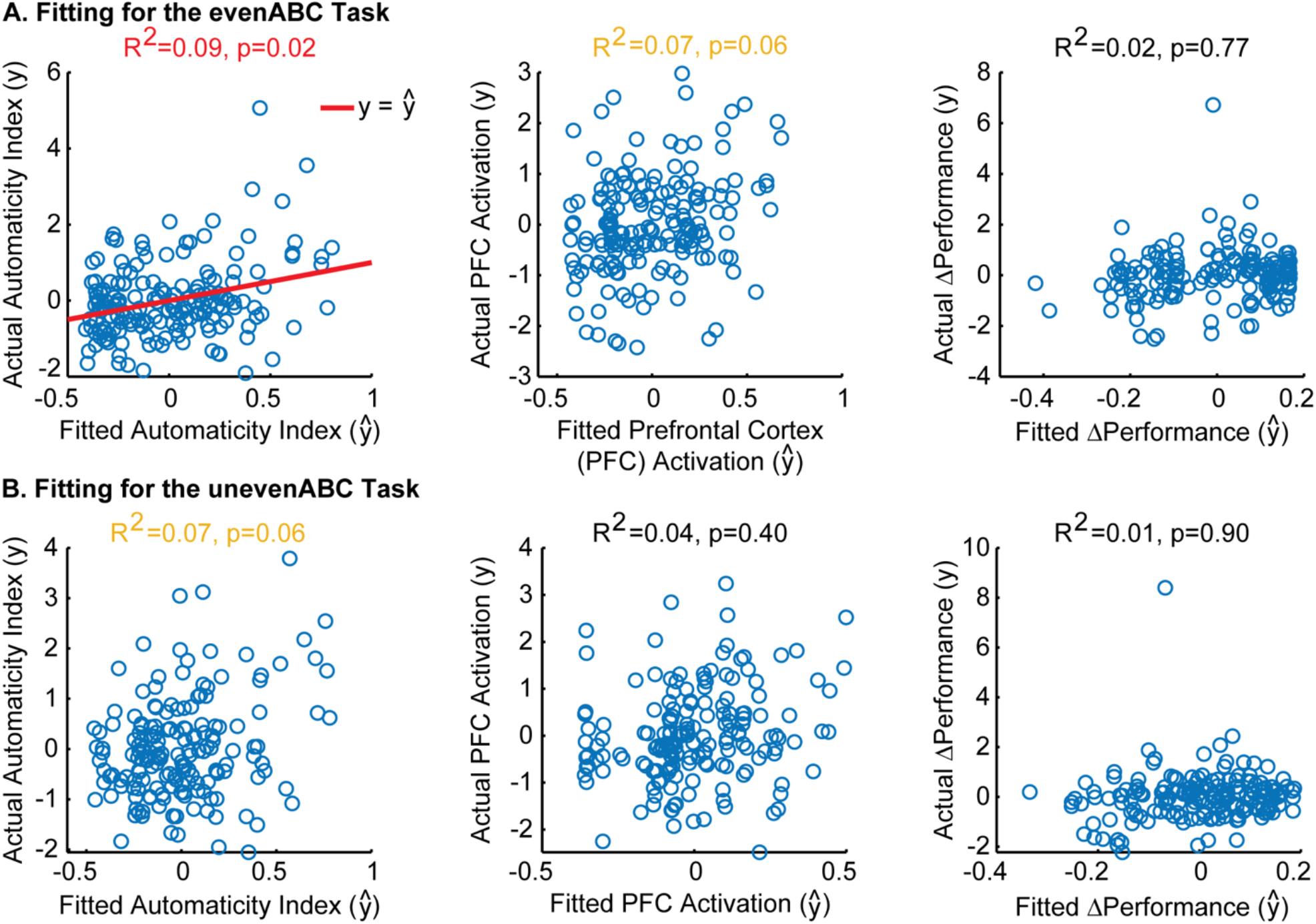
Model fitting between MMSE scores and automaticity index, PFC_Activation_, and Δperformance after adjusting for age, sex, race, and highest level of education. Actual vs. fitted value for each model in the evenABC task **(A)** and unevenABC task **(B)** are shown. Significant models are shown with a red line for *y* = ŷp. Trending models are shown in yellow text. A perfect-fitting model would have actual and fitted values following the *y* = ŷp line. Only the associations between MMSE and automaticity index in the easy task difficulty was significant.

The association between MMSE and gait automaticity had the largest variance accounted for and largest standardized *β* magnitude across both tasks in the unadjusted (Table 2; evenABC: *β* = −0.23 ± 0.07, *p* = 0.002; unevenABC: *β* = −0.22 ± 0.07, *p* = 0.003) and adjusted (Table 3; evenABC: *β* = −0.20 ± 0.08, *p* = 0.01; unevenABC: *β* = −0.19 ± 0.08, *p* = 0.02) models.

The result remains robust after removing the two participants who did not follow instructions to recite every other letter of the alphabet. The models between MMSE and automaticity index still had the largest variance accounted for and largest standardized *β* magnitude across both tasks (Task evenABC: AutomaticityIndex R^2^ = 0.10, *β* = −0.21, PFCActivation: R^2^ = 0.07, *β* = −0.13, ΔPerformance: R^2^ = 0.03, *β* = 0.13. Task unevenABC: AutomaticityIndex R^2^ = 0.08, *β* = −0.20, PFCActivation: R^2^ = 0.04, *β* = −0.10, ΔPerformance: R^2^ = 0.05, *β* = 0.13).

In sum, MMSE score was more strongly correlated with automaticity index than with PFC activation or performance. The association between MMSE scores and automaticity index had the largest standardized *β* magnitude and variance explained across both DT difficulties. The result is robust to adjustments for covariates and not sensitive to the removal of participants who did not follow instructions.

## 4 Discussion

We combined measures of PFC activation and DT performance to create a monotonic index to quantify gait automaticity. We showed that the automaticity index better captured the task difficulty related change in automaticity and was more strongly related to general cognitive function than either PFC activation or DT performance alone. In summary, the proposed index was effective at differentiating 1) between tasks, and 2) between participants.

### 4.1 Task performance contributes valuable information to the automaticity index when comparing task difficulties

When comparing task difficulties, we expected the harder task to show worse automaticity; specifically, the more challenging task will require increased attentional demand from the PFC and result in decreased performance due to greater interference between the motor and cognitive tasks. This expected change would be reflected as an increase in automaticity index, an increase in PFC activation, and a decrease in performance. We observed that 73% of participants increased automaticity index and 71% decreased performance as expected, but only 57% of participants increased PFC activation from evenABC to unevenABC. The result confirms the unevenABC task was more challenging to the participants, given the majority of the participants decreased performance. However, this increased challenge was not always reflected in PFC activation alone.

Multiple reasons could explain why participants did not increase PFC activation with increasing task demands: inefficient recruitment of necessary neural resources, the task exceeding capacity (Reuter-Lorenz and Cappell, 2008), or disengagement from the task such that the participant didn’t even try to cope with the difficulty. In addition to individual differences, the inconsistency in the PFC activity could also be attributed to measurement noise from fNIRS such as physiological changes and skin properties (Vitorio et al., 2017; Menant et al., 2020). This observation emphasizes the heterogeneity of PFC response despite consistent behavioral performance and the importance of incorporating behavioral performance when interpreting PFC activity.

Our finding that DT performance contains crucial information when assessing automaticity aligns with previous studies showing that DT performance is related to older adults’ mobility (Montero-Odasso et al., 2012; Verghese et al., 2012; Rosso et al., 2019a) and cognitive abilities (Holtzer et al., 2006; Rosso et al., 2019b; Brauner et al., 2021). Nevertheless, performance alone does not quantify automaticity since automaticity is defined by proficient performance alongside minimal neural inputs from the attentional and executive control center (Clark, 2015). In other words, the engagement of attentional resources during a typically automatic task such as walking is a signature of reduced automaticity (Clark, 2015; Van Swearingen & Studenski, 2014). Moreover, most theoretical models emphasize the importance of examining the activation patterns of PFC to understand cognitive aging (Reuter-Lorenz and Cappell, 2008; Park and Reuter-Lorenz, 2009; Festini et al., 2018). Considering fNIRS is a relatively new technology with fast-evolving instrument design and analysis techniques (Perrey, 2014), future studies should keep considering performance and PFC activity together using the automaticity index to improve quantification of gait automaticity.

### 4.2 Poorer automaticity index was related to lower general cognitive function

When comparing across participants, we showed that lower cognitive function, measured by MMSE score, is related to lower automaticity index in both DT difficulties. The results suggest that the automaticity index was more sensitive to individual characteristics that could impact automaticity than PFC activation or performance, which are existing metrics to quantify automaticity.

However, we noticed that the variance explained by the models (R^2^) is relatively low. The fit remains roughly the same with or without adjusting for age, sex, and highest level of education, suggesting that most of the variance in the automaticity index and PFC activity was not explained by the demographics and cognitive abilities. The low R^2^ is not unexpected as gait automaticity depends on the subcortical circuits, which were not measured with fNIRS during walking (Wu and Hallett, 2005; Rosano et al., 2007; Clark, 2015). Thus, complementary imaging data about the integrity of the subcortical circuits, including volume or circuitry connectivity, might account for the unexplained variance in the model.

In our study, MMSE was not related to ΔPerformance. In comparison, a prior study (Brauner et al., 2021) has found that MMSE score is associated with ΔPerformance. Of note, ΔPerformance is a component in calculating the automaticity index, but we were not able to observe a direct association between MMSE score and ΔPerformance in our study. Several factors could contribute to this difference, including 1) a different dual-task paradigm was used and 2) the population recruited might be different. Specifically, the prior study had an older cohort than we did (≥ 85 years old vs. mean 73.93 ± 5.65 in our sample).

### 4.3 Automaticity index provides a unified measure that combines behavioral performance and PFC activation

Performance in DT walking is a commonly used metric to assess gait automaticity, but performance alone doesn’t reflect the cortical input required. For instance, the same performance can be achieved with little effort, i.e., automaticity, or with significant effort from the attentional resource center. Neurophysiological measures, such as PFC activation, provide more insights into the cortical inputs supporting the observed performance. However, prior research on PFC signals during walking has shown inconsistent results (Yeung and Chan, 2021). Both increased (Holtzer et al., 2011; Mirelman et al., 2017) and decreased (Beurskens et al., 2014) PFC activity during DT walking have been reported. These findings are hard to reconcile without considering task performance.

The automaticity index provides a solution by establishing a unified axis that combines performance and PFC activation. The increased PFC activity observed by Holtzer et al., 2011 and Mirelman et al., 2017 could indicate either increased reliance on the attentional center to compensate for the loss of automaticity, resulting in maintained task performance, or dedifferentiation in the neural signals, leading to poor task performance. In the automaticity index, the increased PFC activity would be scaled down for maintained performance to reflect successful compensation and scaled up for poor task performance to reflect unsuccessful compensation or de-differentiation. The decreased PFC activity reported by Beurskens et al. (2014) could represent either inefficiency of resource utilization, leading to poor task performance, or maintained automaticity, resulting in good performance. These differences would be reflected in the automaticity index where the same PFC activity would be scaled up for the reduced performance scenario to reflect the inability to recruit necessary resources.

## 5 Limitations

One main limitation is that the automaticity index can only be computed for study designs with at least two distinct task difficulties because parameter optimization *α* requires at least two difficulties. Additionally, even though the proposed method to compute the automaticity index can be applied to any dual-task paradigm with at least two levels of task difficulties, the datasets used here to test the index all performed the same dual-task walking paradigm (i.e., walking and reciting alternating letters of the alphabet with or without an uneven surface). It has been shown that dual-task modality can impact performance and assessment results differently (Beurskens et al., 2014; Tsang et al., 2022). Therefore, it is important to verify that the metric is robust to varying task designs in future studies. Lastly, the participants recruited are relatively healthy in their cognitive abilities (MMSE mean ± SD: 28.42 ± 1.83, min = 21), which may not reflect the full population of community-dwelling older adults (Jacqmin-Gadda et al., 1997). Future studies can assess if the findings are generalizable to populations with a wider range of cognitive abilities.

## 6 Conclusions

Gait automaticity is crucial for safe community mobility and automaticity is an important rehabilitation target to restore walking function and independence. However, a standard and robust assessment for gait automaticity is lacking. We addressed the need to better quantify gait automaticity using a novel approach to combine both DT performance and PFC activation into an automaticity index. We demonstrated the efficacy of the index by achieving two objectives; specifically, better differentiating differences in automaticity 1) between tasks within participants, and 2) between participants based on overall cognitive function. The index revealed a decrease in automaticity as task difficulty increased which was not evidenced by PFC activation. Furthermore, the index captured the cognitive ability-related differences in automaticity better than PFC activation or DT performance alone. In summary, the automaticity index provided a standard metric to characterize gait automaticity in a dual-task walking paradigm. The standardized metric will allow better quantification of the effectiveness of interventions aimed at improving automaticity and facilitate future studies to investigate the connections between automaticity and other participant-specific characteristics.

## Competing interests

The authors have no conflicts of interest to report.

## Author Contributions

SL was involved with study conception, design of the index equations, data analysis and interpretation, and drafting of the manuscript. AR was involved with study conception, data interpretation, and revision of the manuscripts, and provided feedback on the rational of the index. EB collaborated in study conception, data curation, data interpretation, and manuscript revision, and provided feedback on the design and rationale of the index. AW collaborated in the data interpretation and manuscript revision and provided feedback on the rationale of the index. CR reviewed and revised the manuscript. GT was involved with the study conception, design of the index, data interpretation, finding supervision, and manuscript revisions. All authors reviewed and approved the final version of the manuscript.

## Funding

This work was supported by NSF Career Award 1847891 and Pittsburgh Pepper Center P30AG024827. The datasets are from studies supported by RF1AG025516, K01 AG053431 (Neural Mechanisms of Community Mobility); R01-AG045252, K24AG057728, R01 AG057671 (Program to Improve Mobility in Aging); and R01AG023651, U01AG061393 (Move Monongahela-Youghiogheny Healthy Aging Team). AW is supported by the National Institutes of Health (K23AG076663). EB is supported by the National Institutes of Health (T32AG021885).

### Acknowledgments

The authors thank Dr. Dulce M. Mariscal, Dr. Krista Fjeld, Marcela Gonzalez-Rubio, Nathan Brantly, and Adwoa A. Awuah for their insightful feedback on the manuscripts. The authors thank Dr. Ted Huppert for his feedback on the statistical analysis. The authors thank the support from the three studies in sharing the data: Neural Mechanisms of Community Mobility, Program to Improve Mobility in Aging, and Move Monongahela-Youghiogheny Healthy Aging Team.

## Data Availability Statement

The raw data supporting the conclusions of the article will be made available by the authors upon request. The source code supporting the analysis and conclusion of the article is available at https://github.com/PittSMLlab/AutomaticityIndex.git.

## Notes

### Competing Interest Statement

The authors have declared no competing interest.

### Summary of Updates

Manuscript updated for clarity.

## References

Aizenstein, H. J., Nebes, R. D., Saxton, J. A., Price, J. C., Mathis, C. A., Tsopelas, N. D., et al. (2008). Frequent Amyloid Deposition Without Significant Cognitive Impairment Among the Elderly. Arch Neurol 65, 1509. doi: 10.1001/archneur.65.11.1509.

Beurskens, R., Helmich, I., Rein, R., and Bock, O. (2014). Age-related changes in prefrontal activity during walking in dual-task situations: A fNIRS study. International Journal of Psychophysiology 92, 122–128. doi: 10.1016/j.ijpsycho.2014.03.005.

Brach, J. S., VanSwearingen, J. M., Gil, A., Nadkarni, N. K., Kriska, A., Cham, R., et al. (2020). Program to improve mobility in aging (PRIMA) study: Methods and rationale of a task-oriented motor learning exercise program. Contemp Clin Trials 89, 105912. doi: 10.1016/j.cct.2019.105912.

Brandler, T. C., Oh-Park, M., Wang, C., Holtzer, R., and Verghese, J. (2012). Walking while talking: Investigation of alternate forms. Gait Posture 35, 164–166. doi: 10.1016/j.gaitpost.2011.08.003.

Brauner, F. O., Balbinot, G., Figueiredo, A. I., Hausen, D. O., Schiavo, A., and Mestriner, R. G. (2021). The Performance Index Identifies Changes Across the Dual Task Timed Up and Go Test Phases and Impacts Task-Cost Estimation in the Oldest-Old. Front Hum Neurosci 15, 1–12. doi: 10.3389/fnhum.2021.720719.

Brustio, P. R., Rabaglietti, E., Formica, S., and Liubicich, M. E. (2018). Dual-task training in older adults: The effect of additional motor tasks on mobility performance. Arch Gerontol Geriatr 75, 119–124. doi: 10.1016/j.archger.2017.12.003.

Cabeza, R. (2002). Hemispheric asymmetry reduction in older adults: The HAROLD model. Psychol Aging 17, 85–100. doi: 10.1037/0882-7974.17.1.85.

Cabeza, R., and Dennis, N. A. (2012). “Frontal Lobes and Aging,” in Principles of Frontal Lobe Function (Oxford University Press), 628–652. doi: 10.1093/med/9780199837755.003.0044.

Clark, D. J. (2015). Automaticity of walking: Functional significance, mechanisms, measurement and rehabilitation strategies. Front Hum Neurosci 9, 1–13. doi: 10.3389/fnhum.2015.00246.

Clark, D. J., Rose, D. K., Ring, S. A., and Porges, E. C. (2014). Utilization of central nervous system resources for preparation and performance of complex walking tasks in older adults. Front Aging Neurosci 6, 1–9. doi: 10.3389/fnagi.2014.00217.

Festini, S. B., Zahodne, L., and Reuter-Lorenz, P. A. (2018). Theoretical Perspectives on Age Differences in Brain Activation: HAROLD, PASA, CRUNCH—How Do They STAC Up? Oxford Research Encyclopedia of Psychology, 1–24. doi: 10.1093/acrefore/9780190236557.013.400.

Fettrow, T., Hupfeld, K., Tays, G., Clark, D. J., Reuter-Lorenz, P. A., and Seidler, R. D. (2021). Brain activity during walking in older adults: Implications for compensatory versus dysfunctional accounts. Neurobiol Aging 105, 349–364. doi: 10.1016/j.neurobiolaging.2021.05.015.

Folstein, M. F., Folstein, S. E., and McHugh, P. R. (1975). “Mini-mental state”. A practical method for grading the cognitive state of patients for the clinician. J Psychiatr Res 12, 189–98. doi: 10.1016/0022-3956(75)90026-6.

Ganguli, M., Chang, C. C. H., Snitz, B. E., Saxton, J. A., Vanderbilt, J., and Lee, C. W. (2010). Prevalence of mild cognitive impairment by multiple classifications: The monongahela-youghiogheny healthy aging team (MYHAt) project. American Journal of Geriatric Psychiatry 18, 674–683. doi: 10.1097/JGP.0b013e3181cdee4f.

Gschwind, Y. J., Bridenbaugh, S. A., and Kressig, R. W. (2010). Gait disorders and falls. GeroPsych: The Journal of Gerontopsychology and Geriatric Psychiatry 23, 21–32. doi: 10.1024/1662-9647/a000004.

Herold, F., Wiegel, P., Scholkmann, F., Thiers, A., Hamacher, D., and Schega, L. (2017). Functional near-infrared spectroscopy in movement science: a systematic review on cortical activity in postural and walking tasks. Neurophotonics 4, 041403. doi: 10.1117/1.nph.4.4.041403.

Holtzer, R., Mahoney, J. R., Izzetoglu, M., Izzetoglu, K., Onaral, B., and Verghese, J. (2011). fNIRS study of walking and walking while talking in young and old individuals. Journals of Gerontology - Series A Biological Sciences and Medical Sciences 66 A, 879–887. doi: 10.1093/gerona/glr068.

Holtzer, R., Verghese, J., Xue, X., and Lipton, R. B. (2006). Cognitive processes related to gait velocity: Results from the Einstein aging study. Neuropsychology 20, 215–223. doi: 10.1037/0894-4105.20.2.215.

Hoppes, C. W., Huppert, T. J., Whitney, S. L., Dunlap, P. M., Disalvio, N. L., Alshebber, K. M., et al. (2020). Changes in Cortical Activation during Dual-Task Walking in Individuals with and without Visual Vertigo. Journal of Neurologic Physical Therapy 44, 156–163. doi: 10.1097/NPT.0000000000000310.

Jacqmin-Gadda, H., Fabrigoule, C., Commenges, D., and Dartigues, J. F. (1997). A 5-year longitudinal study of the mini-mental state examination in normal aging. Am J Epidemiol 145, 498–506. doi: 10.1093/oxfordjournals.aje.a009137.

Laguë-Beauvais, M., Fraser, S. A., Desjardins-Crépeau, L., Castonguay, N., Desjardins, M., Lesage, F., et al. (2015). Shedding light on the effect of priority instructions during dual-task performance in younger and older adults: A fNIRS study. Brain Cogn 98, 1–14. doi: 10.1016/j.bandc.2015.05.001.

Leys, C., Ley, C., Klein, O., Bernard, P., and Licata, L. (2013). Detecting outliers: Do not use standard deviation around the mean, use absolute deviation around the median. J Exp Soc Psychol 49, 764–766. doi: 10.1016/j.jesp.2013.03.013.

Longhurst, J. K., Rider, J. V., Cummings, J. L., John, S. E., Poston, B., Held Bradford, E. C., et al. (2022). A Novel Way of Measuring Dual-Task Interference: The Reliability and Construct Validity of the Dual-Task Effect Battery in Neurodegenerative Disease. Neurorehabil Neural Repair 36, 346–359. doi: 10.1177/15459683221088864.

Maillet, D., and Rajah, M. N. (2013). Association between prefrontal activity and volume change in prefrontal and medial temporal lobes in aging and dementia: A review. Ageing Res Rev 12, 479–489. doi: 10.1016/j.arr.2012.11.001.

Menant, J. C., Maidan, I., Alcock, L., Al-Yahya, E., Cerasa, A., Clark, D. J., et al. (2020). A consensus guide to using functional near-infrared spectroscopy in posture and gait research. Gait Posture 82, 254–265. doi: 10.1016/j.gaitpost.2020.09.012.

Mirelman, A., Maidan, I., Bernad-Elazari, H., Shustack, S., Giladi, N., and Hausdorff, J. M. (2017). Effects of aging on prefrontal brain activation during challenging walking conditions. Brain Cogn 115, 41–46. doi: 10.1016/j.bandc.2017.04.002.

Miyai, I., Tanabe, H. C., Sase, I., Eda, H., Oda, I., Konishi, I., et al. (2001). Cortical mapping of gait in humans: A near-infrared spectroscopic topography study. Neuroimage 14, 1186–1192. doi: 10.1006/nimg.2001.0905.

Montero-Odasso, M., Muir, S. W., Hall, M., Doherty, T. J., Kloseck, M., Beauchet, O., et al. (2011). Gait variability is associated with frailty in community-dwelling older adults. Journals of Gerontology - Series A Biological Sciences and Medical Sciences 66 A, 568–576. doi: 10.1093/gerona/glr007.

Montero-Odasso, M., Verghese, J., Beauchet, O., and Hausdorff, J. M. (2012). Gait and cognition: A complementary approach to understanding brain function and the risk of falling. J Am Geriatr Soc 60, 2127–2136. doi: 10.1111/j.1532-5415.2012.04209.x.

Park, D. C., and Reuter-Lorenz, P. (2009). The adaptive brain: Aging and neurocognitive scaffolding. Annu Rev Psychol 60, 173–196. doi: 10.1146/annurev.psych.59.103006.093656.

Patla, A. E., and Shumway-Cook, A. (1999). Dimensions of Mobility: Defining the Complexity and Difficulty Associated with Community Mobility. J Aging Phys Act 7, 7–19. doi: 10.1123/japa.7.1.7.

Paul, S. S., Ada, L., and Canning, C. G. (2005). Automaticity of walking–implications for physiotherapy practice. Physical Therapy Reviews 10, 15–23. doi: 10.1179/108331905X43463.

Perrey, S. (2014). Possibilities for examining the neural control of gait in humans with fNIRS. Front Physiol 5 MAY, 10–13. doi: 10.3389/fphys.2014.00204.

Rantakokko, M., Portegijs, E., Viljanen, A., Iwarsson, S., Kauppinen, M., and Rantanen, T. (2016). Changes in life-space mobility and quality of life among community-dwelling older people: a 2-year follow-up study. Quality of Life Research 25, 1189–1197. doi: 10.1007/s11136-015-1137-x.

Reuter-Lorenz, P. A., and Cappell, K. A. (2008). Neurocognitive aging and the compensation hypothesis. Curr Dir Psychol Sci 17, 177–182. doi: 10.1111/j.1467-8721.2008.00570.x.

Rosano, C., Aizenstein, H. J., Studenski, S., and Newman, A. B. (2007). A regions-of-interest volumetric analysis of mobility limitations in community-dwelling older adults. Journals of Gerontology - Series A Biological Sciences and Medical Sciences 62, 1048–1055. doi: 10.1093/gerona/62.9.1048.

Rosso, A. L., Metti, A. L., Faulkner, K., Brach, J. S., Studenski, S. A., Redfern, M., et al. (2019a). Associations of Usual Pace and Complex Task Gait Speeds With Incident Mobility Disability. J Am Geriatr Soc 67, 2072–2076. doi: 10.1111/jgs.16049.

Rosso, A. L., Metti, A. L., Faulkner, K., Redfern, M., Yaffe, K., Launer, L., et al. (2019b). Complex Walking Tasks and Risk for Cognitive Decline in High Functioning Older Adults. Journal of Alzheimer’s Disease 71, S65–S73. doi: 10.3233/JAD-181140.

Rosso, A. L., Taylor, J. A., Tabb, L. P., and Michael, Y. L. (2013). Mobility, disability, and social engagement in older adults. J Aging Health 25, 617–637. doi: 10.1177/0898264313482489.

Santosa, H., Zhai, X., Fishburn, F., and Huppert, T. (2018). The NIRS Brain AnalyzIR toolbox. Algorithms 11. doi: 10.3390/A11050073.

Sheppard, K. D., Sawyer, P., Ritchie, C. S., Allman, R. M., and Brown, C. J. (2013). Life-space mobility predicts nursing home admission over 6 years. J Aging Health 25, 907–920. doi: 10.1177/0898264313497507.

Thies, S. B., Richardson, J. K., DeMott, T., and Ashton-Miller, J. A. (2005). Influence of an irregular surface and low light on the step variability of patients with peripheral neuropathy during level gait. Gait Posture 22, 40–45. doi: 10.1016/j.gaitpost.2004.06.006.

Tsang, C. S. L., Wang, S., Miller, T., and Pang, M. Y. C. (2022). Degree and pattern of dual-task interference during walking vary with component tasks in people after stroke: a systematic review. J Physiother 68, 26–36. doi: 10.1016/j.jphys.2021.12.009.

Van Swearingen, J. M., and Studenski, S. A. (2014). Aging, motor skill, and the energy cost of walking: Implications for the prevention and treatment of mobility decline in older persons. Journals of Gerontology - Series A Biological Sciences and Medical Sciences 69, 1429–1436. doi: 10.1093/gerona/glu153.

Verghese, J., Holtzer, R., Lipton, R. B., and Wang, C. (2012). Mobility stress test approach to predicting frailty, disability, and mortality in high-functioning older adults. J Am Geriatr Soc 60, 1901–1905. doi: 10.1111/j.1532-5415.2012.04145.x.

Verghese, J., Kuslansky, G., Holtzer, R., Katz, M., Xue, X., Buschke, H., et al. (2007). Walking While Talking: Effect of Task Prioritization in the Elderly. Arch Phys Med Rehabil 88, 50–53. doi: 10.1016/j.apmr.2006.10.007.

Vitorio, R., Stuart, S., Rochester, L., Alcock, L., and Pantall, A. (2017). fNIRS response during walking — Artefact or cortical activity? A systematic review. Neurosci Biobehav Rev 83, 160–172. doi: 10.1016/j.neubiorev.2017.10.002.

Wu, T., and Hallett, M. (2005). The influence of normal human ageing on automatic movements. J Physiol 562, 605–615. doi: 10.1113/jphysiol.2004.076042.

Yeung, M. K., and Chan, A. S. (2021). A Systematic Review of the Application of Functional Near-InfraredSpectroscopy to the Study of Cerebral Hemodynamics in Healthy Aging. Neuropsychol Rev 31, 139–166. doi: 10.1007/s11065-020-09455-3.

Yogev-Seligmann, G., Rotem-Galili, Y., Mirelman, A., Dickstein, R., Giladi, N., and Hausdorff, J. M. (2010). How does explicit prioritization alter walking during dual-task performance? Effects of age and sex on gait speed and variability. Phys Ther 90, 177–186. doi: 10.2522/ptj.20090043.

